# Defining how oncogenic and developmental mutations of *PIK3R1* alter the regulation of class IA phosphoinositide 3-kinases

**DOI:** 10.1101/704106

**Authors:** Gillian L. Dornan, Jordan T.B. Stariha, Manoj K. Rathinaswamy, Cameron J. Powell, Martin J. Boulanger, John E. Burke

## Abstract

The class IA PI3Ks are key signalling enzymes composed of a heterodimer of a p110 catalytic subunit and a p85 regulatory subunit, with PI3K mutations being causative of multiple human diseases including cancer, primary immunodeficiencies, and developmental disorders. Mutations in the p85α regulatory subunit encoded by *PIK3R1* can both activate PI3K through oncogenic truncations in the iSH2 domain, or inhibit PI3K through developmental disorder mutations in the cSH2 domain. Using a combined biochemical and hydrogen deuterium exchange mass spectrometry approach we have defined the molecular basis for how these mutations activate both the p110α/p110δ catalytic subunits. We find that the oncogenic Q572* truncation of *PIK3R1* disrupts all inhibitory inputs, with p110α being hyper-activated compared to p110δ. In addition, we find that the R649W mutation in the cSH2 of *PIK3R1* decreases sensitivity to activation by receptor tyrosine kinases. This work reveals novel insight into isoform specific regulation of p110s by p85α.

**Highlights:**

- An oncogenic variant of p85α, Q572*, leads to hyper-activation of p110α compared to p110δ
- HDX-MS revealed that Q572* leads to disruption of all inhibitory interfaces in p110α
- A SHORT syndrome mutation in p85α leads to decreased sensitivity to RTKs for p110α/δ

## Introduction

The class I phosphoinositide 3-kinases (PI3Ks) are lipid signalling enzymes that generate the lipid signal phosphatidylinositol (3,4,5) trisphosphate (PIP3) downstream of cell surface receptors, including receptor tyrosine kinases (RTKs), G-protein coupled receptors (GPCRs), and Ras (Burke, 2018; Burke and Williams, 2015). Generation of PIP3 leads to the activation of many pro-growth signalling events, including activation of the protein kinase Akt, resulting in increased nutrient uptake, cell growth, cell proliferation, and cell survival (Fruman et al., 2017). PI3Ks are able to respond downstream of diverse RTKs in different cell types including epidermal growth factor receptors (EGFR), platelet derived growth factor receptor, and the insulin receptor. The class I PI3Ks can be split into two distinct subgroups (class IA and IB) dependent on their ability to be activated downstream of distinct cell surface receptors and through engagement of different regulatory proteins. Misregulated PI3Ks play critical roles in numerous human diseases, which are mediated by both upregulated and downregulated PI3K activity. Activating mutations of class IA PI3Ks occur frequently in cancer (Samuels et al., 2004), primary immunodeficiencies (Angulo et al., 2013; Lucas et al., 2016; 2014a), and overgrowth developmental disorders (Lindhurst et al., 2012; Venot et al., 2018). Intriguingly, inactivating mutations in class IA PI3Ks are also found in immune disorders (Sharfe et al., 2017), and developmental disorders (Chudasama et al., 2013; Dyment et al., 2013; Thauvin-Robinet et al., 2013), highlighting the critical role of properly regulating PI3K activity.

The class IA PI3K heterodimers are composed of three distinct catalytic subunits (p110α, p110β, or p110δ) and one of five distinct regulatory subunits (p85α, p55α, p50α, p85β, or p55γ), with the regulation of p110 lipid kinase activity being critically dependent on the formation of a heterodimeric complex. The regulatory subunit plays three key roles in regulating p110 subunits: stabilise the p110 subunit, inhibit basal lipid kinase activity, and allow for activation downstream of pYXXM motifs in phosphorylated receptors and their adaptors (Yu et al., 1998a; 1998b). Both p110 and p85 subunits are large multi-domain proteins. The p110 subunits are composed of an adaptor binding domain (ABD) that mediates binding to the iSH2 of p85, a Ras binding domain (RBD) that mediates activation downstream of Ras (Pacold et al., 2000), a C2 domain, a helical domain and a bi-lobal kinase domain (Huang et al., 2007). The five regulatory subunits all contain three essential regulatory domains, the nSH2 and cSH2 domains that mediate binding and activation downstream of pYXXM motifs (Piccione et al., 1993; Rordorf-Nikolic et al., 1995; Songyang et al., 1993) and an interspersing iSH2 coiled coil domain that mediates the high affinity interaction with p110 subunits (Miled et al., 2007). Biophysical and biochemical studies have defined a set of inter and intra subunit inhibitory interactions that mediate p110 inhibition. These four regulatory interfaces are a p110 intra-subunit ABD-kinase interface, an inter-subunit C2-iSH2 interface, an inter-subunit interaction between the nSH2 and the C2, helical, and kinase domains, and an inter-subunit interaction between the cSH2 and the kinase domain (only present for p110β and p110δ) (Vadas et al., 2011).

The different p110 isoforms are uniquely regulated by the different domains of p85, with the nSH2 domain mediating inhibition of all class IA p110 subunits through direct interaction with the C2, helical and kinase domains (Mandelker et al., 2009; Miled et al., 2007), however, the cSH2 only forms an inhibitory interaction with the kinase domain of p110β and p110δ (Burke et al., 2011; Zhang et al., 2011). The nSH2 domain interaction with p110 occludes the phosphotyrosine binding site, as this site is directly involved in the interaction with p110 (Mandelker et al., 2009). The iSH2 domain inhibits all three p110 subunits through an interaction with their C2 domains, though this interaction is partially disrupted in p110β (Burke and Williams, 2013; Dbouk et al., 2010). The iSH2 is composed of three α helices (α1 (441-513), α2 (515-588), α3 (590-599)) with two extensive α helices (α1+α2) that mediate binding to p110 subunits, and a short α3 helix that is located in close spatial proximity to the activation loop of p110. The α3 helix of the iSH2 is dynamic when in a complex with p110 catalytic subunit, as electron density for this helix is not well defined in all of the multiple crystal structures of different p110-iSH2 complexes.

The p85α regulatory subunit (encoded by the *PIK3R1* gene) is the most frequently mutated PI3K regulatory subunit in human disease (Dornan and Burke, 2018; Nunes-Santos et al., 2019), including activating mutations that mediate oncogenic transformation and primary immunodeficiencies, and inactivating mutations that disrupt activation downstream of SH2 interactions with phosphorylated receptors and adaptors. Furthermore, loss of p85α leads to oncogenic transformation mediated by p110α, suggesting that this specific regulatory subunit acts as a tumour suppressor (Thorpe et al., 2017). The first identified oncogenic mutation of *PIK3R1* results in premature truncation at Q572, located within α2 of the iSH2 domain, leading to constitutive activation of PI3K (Jaiswal et al., 2009; Jimenez et al., 1998; Shekar et al., 2005). Multiple additional activating oncogenic point mutations/truncations of *PIK3R1* have been identified, primarily localised at the C2-iSH2 interface (Jaiswal et al., 2009; Sun et al., 2010; Wu et al., 2009), with oncogenic transformation mediated by p110α (Sun et al., 2010). Activating splice mutations in *PIK3R1* lead to primary immunodeficiencies, a condition known as activated PI3K delta syndrome 2 (APDS2), resulting from a truncated p85α protein missing the n-terminus α1 of the iSH2 domain (Δ434-475) (Deau et al., 2014; Lucas et al., 2016; 2014b). Intriguingly this mutation leads to selective activation of p110δ over p110α, providing a putative mechanism for why this mutation primarily leads to a p110δ mediated primary immunodeficiency (Dornan et al., 2017). Finally, mutations in *PIK3R1* are a frequent cause of a developmental disorder (SHORT syndrome) (Chudasama et al., 2013; Dyment et al., 2013; Thauvin-Robinet et al., 2013) mediated by truncations, insertions, or point mutations in the cSH2 domain of p85α that disrupt binding to pYXXM motifs in RTKs.

Recently, we identified unexpected, isoform-specific differences in the regulation of class IA PI3K p110 isoforms by activating p85 mutations (Dornan et al., 2017). Thus, we set out to investigate the molecular mechanisms by which oncogenic and developmental disorder mutations in p85α result in aberrant regulation of p110. Using a combined biochemical and biophysical approach, we first characterised the effects of oncogenic truncations in the iSH2 of p85α on activation of all p110 isoforms. We then investigated the R649W mutation in the cSH2 of p85α, which causes SHORT syndrome, and defined the mechanism by which this mutation leads to disruption of activation downstream of phosphorylated receptors and their adaptors. The conformational changes that occur upon mutation of p85α, and the resulting effects on activation downstream of RTKs, were probed using hydrogen deuterium exchange mass spectrometry (HDX-MS). We find that oncogenic truncations (Q572*) of *PIK3R1* fully disrupt SH2-mediated activation of all p110 catalytic isoforms, while SHORT mutations in the cSH2 of *PIK3R1* (R649W) lead to a disruption in the ability of p110 isoforms to be activated by bis-phosphorylated peptides derived from RTKs. Overall, this work provides unique insight into the molecular mechanisms mediating PI3K inhibition and activation.

## Results

We previously defined the molecular mechanism of APDS2 splice mutations of *PIK3R1* that lead to deletion of the N-terminal region of the α1 helix of the iSH2, and found that it specifically activates the p110δ isoform over the p110α catalytic isoform (Dornan et al., 2017). We wanted to understand how other deletions and truncations of the iSH2 domain affect the regulation of different p110 catalytic isoforms. To this end, we produced oncogenic and engineered deletions of the iSH2 domain of p85α to probe the role of the iSH2 and the cSH2 domains in the regulation of both p110δ and p110α (Fig 1A). We generated two oncogenic truncations of p85α (Q572* (Jimenez et al., 1998) and E601* (Giannakis et al., 2016), that lead to a loss of part of the α2 helix onward or loss of just the cSH2 respectively, and an engineered truncation that removes the α3 helix of the iSH2 (R590*). These truncations were made in complex with both p110δ and p110α from baculovirus infected Sf9 cells (Fig. 1A-C). All constructs expressed well, and eluted from gel filtration at a volume consistent with them being a heterodimer of the p110 and p85 subunits (Fig. S1). These experiments allowed us to probe the role of the C-terminus of the iSH2 domain, as well as the cSH2 domain in the regulation of both p110δ and p110α.

**Figure 1.**
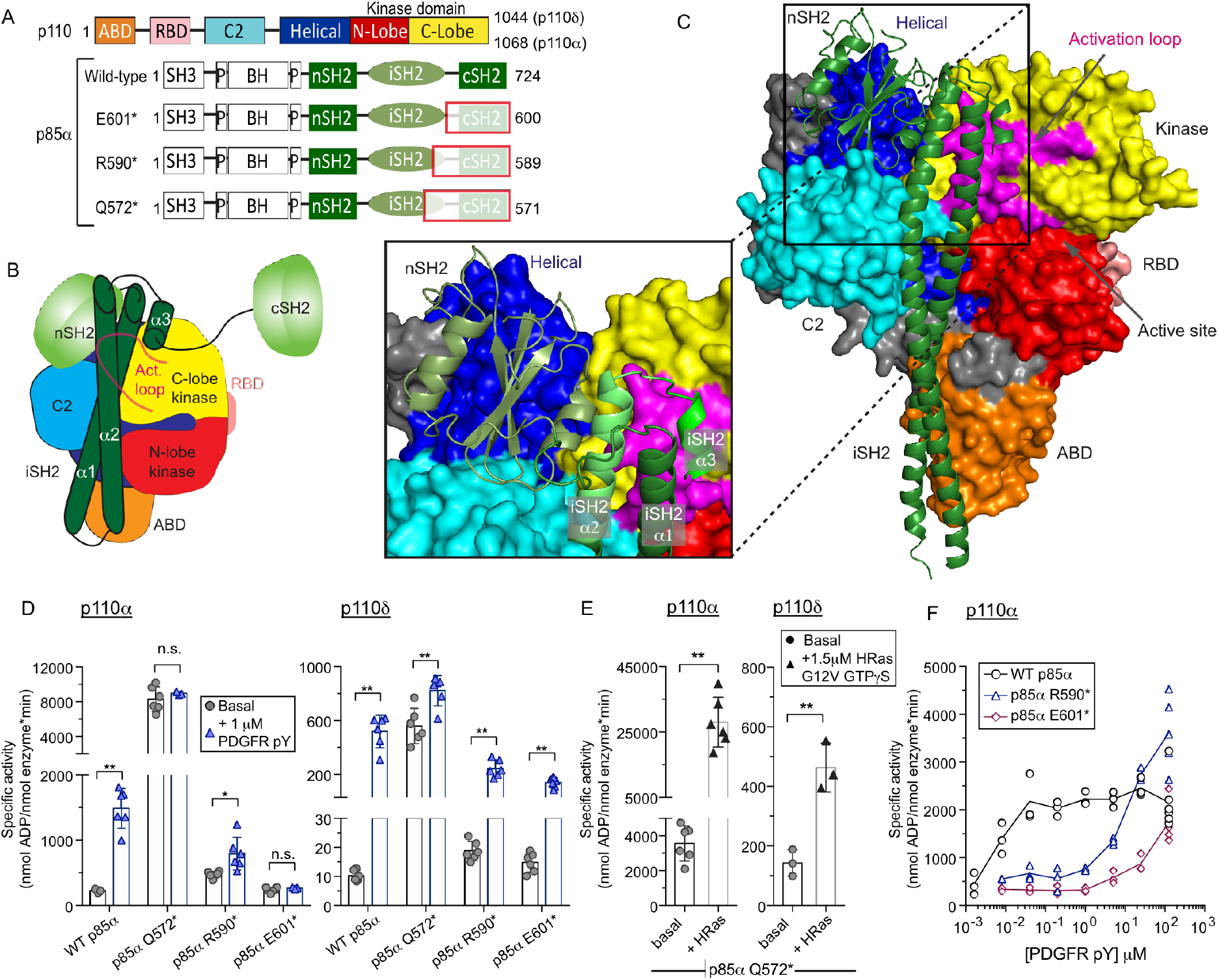
C-terminal truncation of p85α leads to decreased sensitivity to PDGFR stimulation. **(A)** Primary sequence of p110α, p110δ, and p85α wild-type and mutant constructs. Truncated regions indicated by white boxes with red outline. See also Figure S1. **(B)** Schematic model demonstrating the domain architecture of PI3Kα (p110α/p85α). The iSH2 domain is composed of three α helices: α1, α2, and α3. **(C)** A structural model of PI3Kα (p110α/p85α) with a zoom-in representation of the three iSH2 helices. The α1 and α2 helices form the major anti-parallel coiled-coil interface and the α1 helix also forms an interface with the nSH2 domain and the kinase domain. The smaller α3 helix forms interfaces with the α1 and α2 helices as well as the activation loop of the kinase domain. **(D)** Lipid kinase activity of the wild-type and C-terminal truncations of PI3Kα (p110α/p85α) and PI3Kδ (p110δ/p85α) in the presence (+ 1 μM PDGFR pY) or absence (Basal) of a stimulating PDGFR derived phosphopeptide. The phosphopeptide corresponds to residues 735-767 of the PDGFR-β kinase insert region, with residues Y740 and Y751 phosphorylated (pY740 and pY751). All PDGFR pY tested conditions are the bis-phosphorylated peptide unless otherwise noted. See also Fig. S2 **(E)** Specific activities of C-terminal truncations PI3Kα (p110α/p85α) and PI3Kδ (p110δ/p85α) on plasma membrane-mimic vesicles in the basal state and in the presence of 1.5 μM of lipidated HRas G12V loaded with GTPγS. **(F)** Dose response of bis-phosphorylated PDGFR phosphopeptide concentration of the wild-type and the C-terminal truncations R590* and E601*. All assays measured the production of ADP in the presence of 0.1– 1100 nM of enzyme, 100 μM ATP, and PM mimic vesicles containing 5% PIP2. Kinase assays were performed in triplicate (error shown as SD; n = 3-6). Differences between conditions tested using the standard students T-test (two-tailed, unequal variance). * = p< 0.05 (95% confidence interval). ** = p< 0.01 (99% confidence interval). n.s. = p > 0.05.

We measured the lipid kinase activity of the C-terminal truncations of p85α in the context of both p110δ and p110α using lipid vesicles mimicking the composition of the plasma membrane (5% PIP2, 30% PS, 50% PE, 15% PC). Lipid kinase activity of the enzymes was measured in the absence (basal) and presence of 1 μM of a PDGFR bis-phosphorylated pY peptide (referred to going forward as PDGFR pY) composed of PDGFR residues 735-767 with phosphorylation present at Y740 and Y751. The presence of PDGFR pY leads to activation of both p110α and p110δ consistent with previous results (Fig. 1D). The oncogenic truncation of p85α (Q572*) activates both p110α and p110δ, with it leading to significantly higher (5-6 fold higher) lipid kinase activity than PDGFR pY activation in p110α with full length p85α (Fig. 1D). Intriguingly, in p110δ Q572* only activates to the same level as PDGFR pY activation. There is no further activation in the presence of PDGFR-pY for the Q572* construct for p110α, with a slight activation with p110δ. This is consistent with previous studies that showed the p85α-Q572* mutation binds but does not inhibit p110α (Shekar et al., 2005; Wu et al., 2009). To determine if either p110α or p110δ bound to Q572* represented a fully activated form we tested the activation by lipidated HRas, which activates p110α and p110δ through increased membrane recruitment. We found that lipidated HRas activated both p110α and p110δ bound to Q572* similar to the activation seen with wildtype p85α (Fig 1E) (Buckles et al., 2017; Siempelkamp et al., 2017).

The other truncations of p85α (R590* and E601*) in complex with p110α showed little (R590*) or no activation (E601*) compared to wild type p85α in the basal state. Intriguingly, for p110α the presence of 1 μM PDGFR-pY caused no activation for the E601* truncation, and only very minor activation in R590*. For p110δ, both R590* and E601* led to weak activation of basal activity compared to WT p85α, consistent with studies on the role of the cSH2 domain in inhibiting lipid kinase activity of p110δ through an interaction with the kinase domain (Burke et al., 2011). In contrast to p110α, PDGFR-pY activation of E601* and R590* bound to p110δ led to weak activation of lipid kinase activity, although not at the same level as the WT full length p85α, revealing that even though the cSH2 domain plays a key inhibitory role for p110δ it is less important for promoting activation downstream of PDGFR-pY (Fig. 1D). To determine whether the ability of p110α to be activated by PDGFR pY was completely abolished with the truncated E601* and R590* constructs, we carried out lipid kinase assays across a large range of PDGFR pY concentrations (1 nM to 125 μM) (Fig. 1E). The response of p110α in the presence of WT p85α showed activation with an EC50 of roughly 5 nM, consistent with previous studies (Burke and Williams, 2013). The E601* construct bound to p110α was able to be fully activated to the same extent as WT p85α, however this only occurred at a greater than 1000-fold increase in concentration of PDGFR-pY. The activation of R590* bound to p110α occurred at a similar concentration, however, this construct was able to be hyper-activated beyond WT p85α, highlighting a role of the α3 helix in partially inhibiting enzyme activity. These data indicate that the cSH2 is required for efficient activation of p110α despite the cSH2 not forming inhibitory interfaces with the catalytic subunit as in other isoforms, and that loss of the cSH2 leads to decreased sensitivity to activation by PDGFR pY, consistent with previous results (Carpenter et al., 1993; Klippel et al., 1992; Piccione et al., 1993; Rordorf-Nikolic et al., 1995). These same experiments carried out with the E601* p85 construct with p110δ showed a similar fold decrease in sensitivity to pY activation, revealing that the cSH2 is likely critical in sensitivity to PDGFR for all class IA isoforms (Fig. S2).

### The p85α oncogenic Q572* truncation of the iSH2 coiled-coil leads to disruption of key inhibitory interfaces

To investigate the molecular mechanisms that lead to the hyper-activation of the p85α-Q572* truncation mutant when in complex with p110α, we used HDX-MS to probe for activating conformational changes between the WT and the p85α-Q572* truncation. HDX-MS is a powerful analytical technique that measures the exchange rate of amide hydrogens in solution, and has been extensively used to define the conformational mechanisms that underlie PI3K activation by both activating stimuli and disease linked mutations (Burke, 2019; Masson et al., 2017; Vadas and Burke, 2015). Essential to this technique is the generation of peptic peptides spanning the entire sequence to allow for the localisation of deuterium incorporation. The full set of all peptides analysed for both p110α and p85α is shown in Source data 1.

HDX-MS experiments were carried out at three time points of deuterium exchange (3, 30 and 300 seconds) for p110α bound to either wild type p85α or the Q572* truncation. There were multiple regions in both the p110α and p85α subunits that showed significant increases in exchange in the Q572* mutant (Fig. 2A+B). These increases were primarily localised at key inter and intra subunit inhibitory interfaces. These regions include p110α peptides spanning the ABD-kinase interface (10-19), the ABD-RBD linker (100-119, 120-127), the C2-iSH2 interface (343-350), the C2-nSH2 interface (444-455) and p85α peptides spanning the nSH2-helical interface (371-380) and the iSH2-C2 interface (467-476 and 556-570). There were large increases in exchange in the p85α-Q572* truncation at all regions of the iSH2 not in contact with the ABD domain. This is in agreement with previous studies that the p85α-Q572* mutation in complex with p110α leads to disruption of the iSH2-C2 interface and explains why there is no further activation in the presence of the N564D mutation present at the C2-iSH2 interface (Wu et al., 2009). Overall the increased H/D exchange at all inhibitory interfaces reveals the molecular mechanism for activation, with the increases in H/D exchange being similar to previously observed increases in exchange that occur in p110α oncogenic variants or upon PDGFR-pY activation and membrane binding (Burke et al., 2012; Burke and Williams, 2013). Overall, these results revealed that the Q572* mutation induces conformational changes that mimic activating oncogenic p110α mutants.

**Figure 2.**
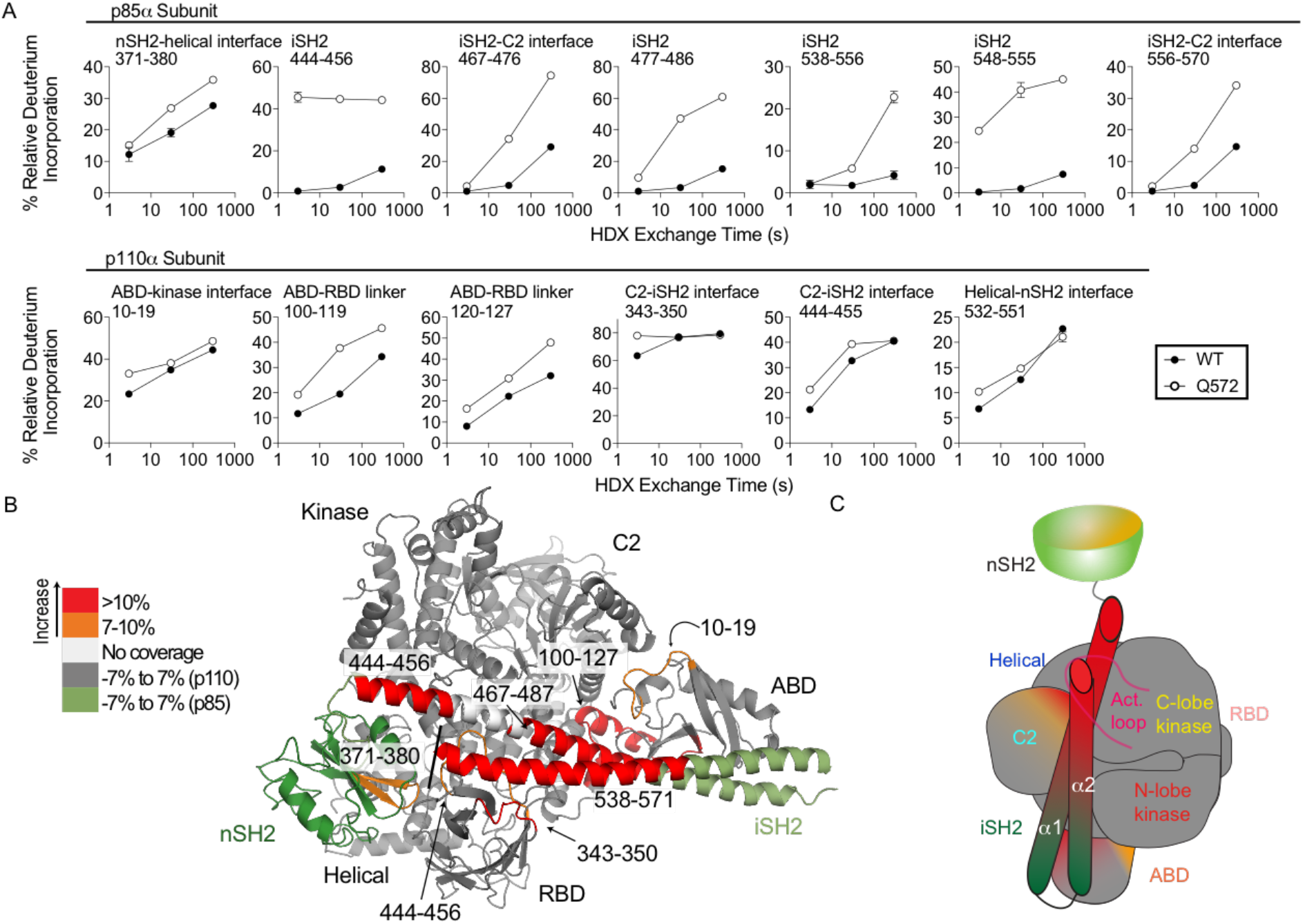
Hydrogen deuterium eXchange reveals disruption of key inhibitory interfaces in the Q572* C-terminal truncation mutant. **(A)** Time course of deuterium incorporation for a selection of peptides in both p85α and p110α with HDX differences in the Q572* mutant. **(B)** Peptides in p85α and p110α that showed differences in HDX that were greater than both 0.4 Da and 7% in the Q572* C-terminal truncation mutant compared with the WT are mapped on the structure of p110α bound to both the nSH2 and iSH2 of p85α [PDB: 2OVU (Miller et al., 2014)]. **(C)** Schematic model of conformational changes that occur in the complex with the Q572* mutation that leads to C-terminal truncation of p85α.

### Both SH2 domains are required for full and efficient PI3K activation

The massive decrease in sensitivity of the p85α E601* construct with p110α to PDGFR pY suggested a key role of the cSH2 in mediating activation of this isoform. We speculated that this construct may act similarly to the previously identified SHORT mutation (R649W) (Chudasama et al., 2013; Dyment et al., 2013; Thauvin-Robinet et al., 2013) that is proposed to disrupt phosphopeptide binding of the cSH2 domain. To study this comprehensively, we generated p85α variants containing either the SHORT mutation in the cSH2 (R649W) or a previously characterised engineered mutation in the nSH2 domain (R358A) (Yu et al., 1998a). Both of these mutations lead to disruption of the FLVR motif in the SH2 domains that mediates binding to phosphopeptide (Fig 3A) (Breeze et al., 1996; Nolte et al., 1996).

We determined the lipid kinase activity of p110α bound to three different p85α constructs (WT, R358A, and R649W) in the presence of the PDGFR pY peptide or PDGFR peptides monophosphorylated at either pY740 or pY751 (Fig. 3B-E). Lipid kinase assays on the R358A and R649W showed no significant activation of basal lipid kinase activity (Fig. 3B). Intriguingly, there was no activation of p110α / R649W p85α in the presence of 1 μM PDGFR-pY similar to p110α / E601*, with a small but significant activation by PDGFR-pY of the p110α / R358A p85α complex. This is surprising as removal of the nSH2 is required for activation through the disruption of the nSH2 inhibitory interaction. To test whether higher concentrations of PDGFR pY would lead to full activation of the mutants, we conducted a similar dose response activity experiment with the full complement of phosphorylated peptides (1.6 nM-125 μM). A PDGFR pY dose response revealed that the SHORT mutation in the cSH2 mimicked the dose response of E601*, where full activation was able to be achieved at high concentrations of PDGFR pY (Fig. 3C). This shows that the nSH2-helical interface is still broken, however the cSH2 binding of pYXXM is required for sensitivity to PDGFR pY. The nSH2 FLVR mutant revealed activation at lower concentrations of PDGFR pY, however, even at high concentrations the nSH2 FLVR mutant was not able to be fully activated compared to WT p85α (Fig. 3C).

**Figure 3.**
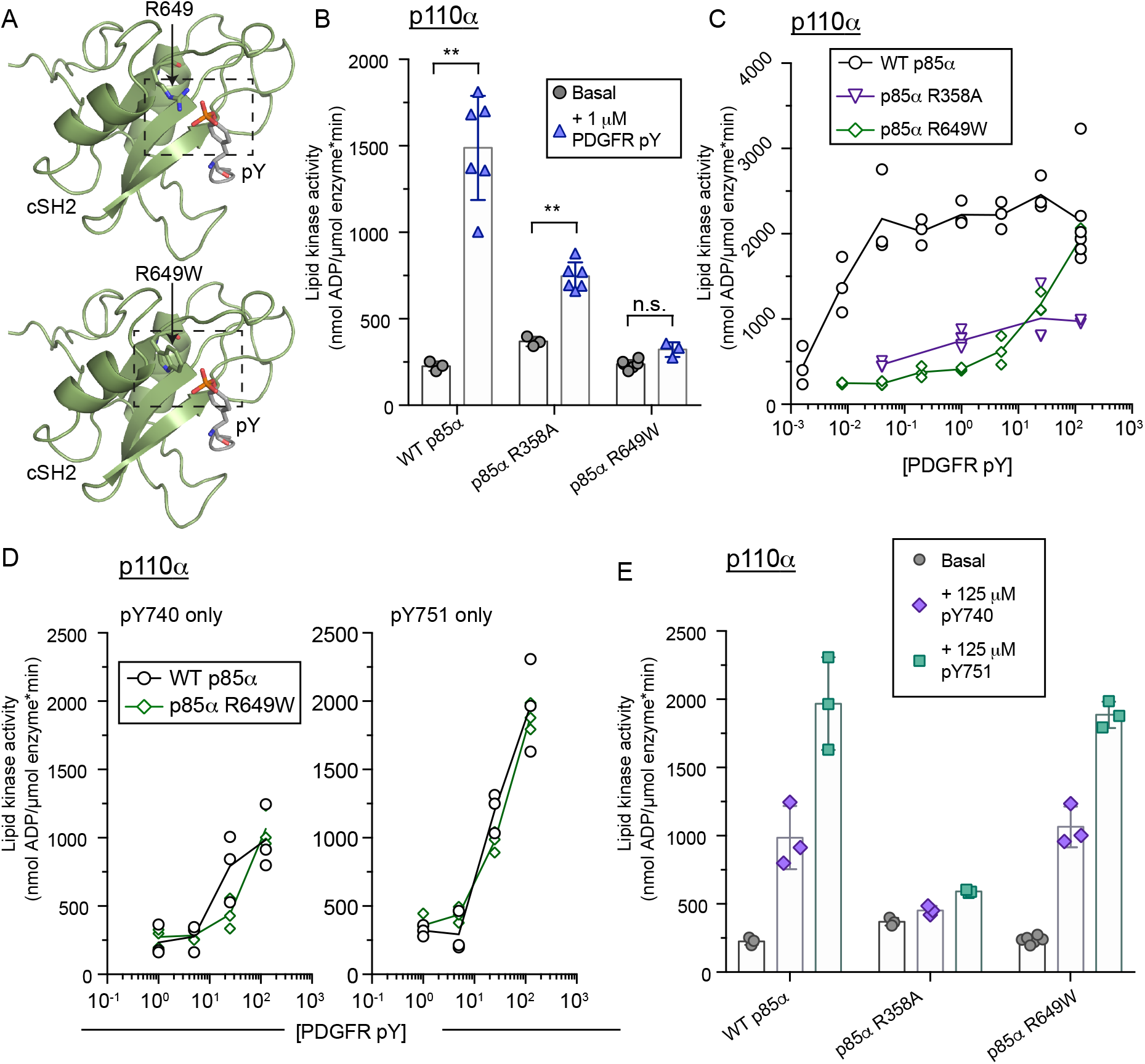
SHORT mutation of key phosphopeptide binding residue leads to decreased phosphopeptide sensitivity. **(A)** Structural model of the R649W mutation in the cSH2 domain of *PIK3R1*. The R649W mutation model was generated in COOT (Emsley et al., 2010) from the cSH2 structure bound to phosphorylated PDGFR (PDB 1PIC) (Hoedemaeker et al., 1999), selecting the highest frequency rotamer of W649 with the lowest clash score. **(B)** Lipid kinase activity of the WT PI3Kα (p110α/p85α) and PI3Kα with single SH2 FLVR mutations (p110α/p85α-R358A and p110α/p85α-R649W) in the absence (Basal) or presence (+1 μM PDGFR pY) of PDGFR bis-phosphorylated pY. Assays measured the production of ADP in the presence of 0.1–100 nM of enzyme, 100 μM ATP, and PM mimic vesicles containing 5% PIP2. Kinase assays were performed in triplicate (error shown as SD; n = 3-6). **(C)** Dose response of bisphosphorylated PDGFR pY concentrations of WT PI3Kα (p110α/p85α) and PI3Kα with single SH2 FLVR mutations (p110α/p85α-R358A and p110α/p85α-R649W). Lipid kinase activity was measured at four to eight different concentrations of PDGFR pY (0.0016 nM – 125 μM). Assays measured the production of ADP in the presence of 0.1–100 nM of enzyme, 100 μM ATP, and PM mimic vesicles containing 5% PIP2. Kinase assays were performed in triplicate (error shown as SD; n = 3). **(D)** Dose response of single phosphorylated PDGFR pY peptides (Peptides with either pY740 or pY751 as indicated above the graph) concentration of the wild-type and the SH2 FLVR mutation R649W. Lipid kinase activity was measured at four different concentrations of PDGFR pY (1 nM – 125 μM). Assays measured the production of ADP in the presence of 0.1–100 nM of enzyme, 100 μM ATP, and PM mimic vesicles containing 5% PIP2. Kinase assays were performed in triplicate (error shown as SD; n = 3). See also Fig. S3. **(E)** Lipid kinase activity of the WT PI3Kα (p110α/p85α) and PI3Kα with single SH2 FLVR mutations (p110α/p85α-R358A and p110α/p85α-R649W) in the absence (Basal) or presence (+125 μM PDGFR pY) of high concentration of PDGFR mono-phosphorylated pY peptides. Assays measured the production of ADP in the presence of 0.1–100 nM of enzyme, 100 μM ATP, and PM mimic vesicles containing 5% PIP2. Kinase assays were performed in triplicate (error shown as SD; n = 3-6).

To determine if the decreased sensitivity to PDGFR-pY activation in R649W p85α was due to the cSH2 providing a template for nSH2 binding of the bis-phosphorylated peptide, we carried out lipid kinase assays in the presence of varying concentrations (1 μM-125 μM) of both single mono-phosphorylated PDGFR pY peptides (Fig. 3D). When incubated with PDGFR pY740, both the p110α bound to p85α WT or p85α R649W showed activation at high concentrations, although it was never able to reach the same activity as with the bis-phosphorylated peptide. This is consistent with previous data showing that the nSH2 has greatly decreased affinity for this site compared to the cSH2 (Panayotou et al., 1993). With the PDGFR pY751 mono-phosphorylated peptide, both the WT and the R649W constructs were able to be fully activated, however, activation required >100 fold more peptide than WT p85α with the bis-phosphorylated phosphopeptide. The R358A construct bound to p110α was only very weakly activated by the mono-phosphorylated peptides (Fig. 3E). Previous experiments showed that both the nSH2 and the cSH2 domains can bind the pY751 site with high affinity (Panayotou et al., 1993), with this likely explaining the increased activation with this peptide over the pY740 peptide. To define the effect of the R649W mutation on binding, we carried out isothermal titration calorimetry experiments using both mono-phosphorylated peptides (Fig. S3 and Table S1). Initially, we measured the affinities between wild type p110α / p85α and the mono-phosphorylated peptides to be in the range of 15-30 nM, consistent with previously determined constants determined for the cSH2 alone (O’Brien et al., 2000; Panayotou et al., 1993). We next measured binding of the complex of p110α / R649W p85α to mono-phosphorylated peptides and showed no detectable binding, in contrast to what we observed in lipid kinase assays. We reasoned that the lack of detectable binding was the result of complications in the measurement due to the combination of phosphopeptide binding to the nSH2 and disruption of the nSH2-p110 interaction.

### Absence of the cSH2 leads to decreases in conformational changes associated with full class IA PI3K activation

To investigate how the C-terminal truncations were mediating the observed PDGFR pY activity differences, we used HDX-MS to examine conformational changes upon pY binding for p110α bound p85α WT, R590*, and R649W. The p110α/p85α constructs were incubated with 0 μM, 1 μM, or 20 μM PDGFR pY and exchange reactions proceeded for three different time points (3, 30, 300s). The full set of all peptides analysed for both p110α and p85α is shown with the level of deuterium incorporation at every time point and condition in the source data.

The WT p110α/p85α heterodimer exhibited changes in deuterium incorporation with 1 μM PDGFR pY consistent with previously published HDX-MS results, with no additional changes occurring with the higher concentration (20 μM) of PDGFR-pY (Fig. 4A). These changes were localised at key regulatory interfaces including the helical-nSH2 interface of p110α (532-551), the C2-nSH2 interface (444-455), the ABD-RBD linker (100-119), and the N and C termini of the p85-iSH2 (444-456 and 582-596) (Fig. 4B+C). Both the nSH2 and the cSH2 domains showed large decreases in exchange consistent with protection due to PDGFR pY binding. Both the R590* and R649W p85α mutant PI3Kα showed very few differences in exchange in the presence of 1 μM pY. However, in the presence of 20 μM pY both of these constructs showed changes in deuterium incorporation similar to the WT p85α at 1 μM pY (Fig. 4B). This indicates that the C-terminal truncations are being activated by the same mechanisms as WT but are less sensitive to the PDGFR pY, consistent with biochemical assays. The p85α-R590* mutant also showed increased exchange compared to WT p85α in both the α1 and α2 helices of the iSH2 (444-456, 556-570), which is the putative contact area of the α3 helix. This is consistent with truncation of the α3 helix leading to a weak activation of PI3K activity both basally and upon addition of PDGFR pY at 1 μM (Fig. 1). If the α3 helix had no regulatory role, we would expect to see a similar kinase activity to the p85-E601* mutant. Interestingly, the cSH2 domain of the p85α-R649W mutant still exhibits decreases in deuterium incorporation in the cSH2 associated with PDGFR pY binding (681-687 and 704-710). These changes occurred only in the presence of 20 μM PDGFR pY, and were smaller than the decreases seen in WT p85α. This suggests that upon engagement of the nSH2 domain by the bis-phosphorylated peptide, that there can still be interaction with the mutated cSH2.

**Figure 4.**
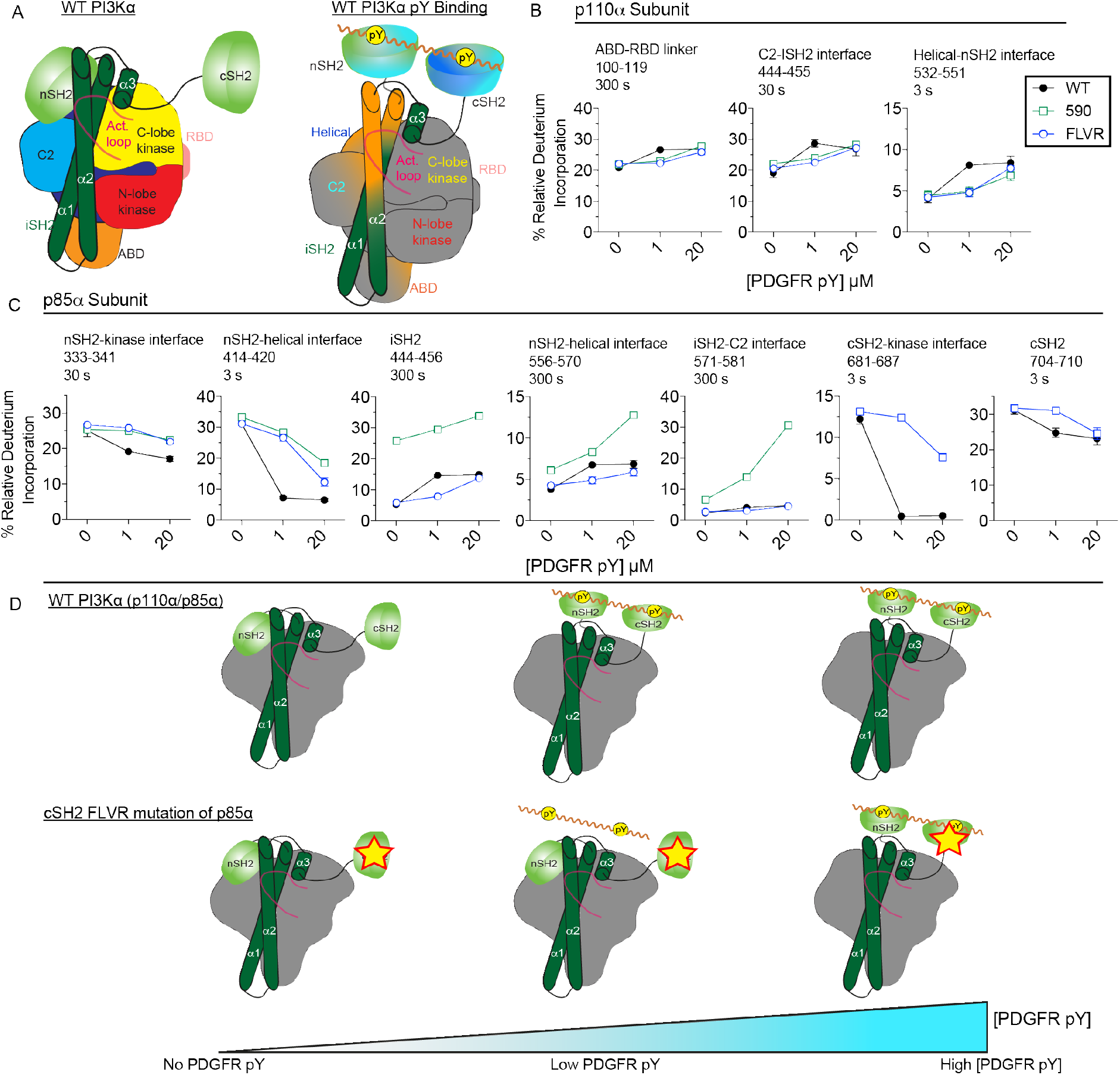
Hydrogen deuterium eXchange reveals decreased sensitivity of C-terminal variants compared to wild-type PI3Kα. **(A)** Peptides in p85α and p110α that showed differences in HDX that were greater than both 0.4 Da and 4% in the wild-type PI3Kα in the absence or presence of PDGFR pY (1 or 20 μM) peptide are mapped on a schematic model of WT PI3Kα (p85α/p110α) to indicate the conformational changes that occur in the WT complex upon binding to 1 μM PDGFR pY peptide. **(B)** The changes in % relative deuterium incorporation of PI3K variants in the presence of different PDGFR pY peptide concentrations (0, 1, or 20 μM). Graphs represent deuterium incorporation of peptides in the p110α of wild-type and p85α variants (WT, R590*, and R649W) at specific time points. The p110α/p85α-E601* variant was also tested but did not undergo changes in deuterium incorporation that met our threshold. **(C)** The changes in % relative deuterium incorporation of PI3K variants in the presence of different PDGFR pY peptide concentrations (0, 1, or 20 μM). Graphs represent deuterium incorporation of peptides in the p85α of wild-type and p85α variants (WT, R590*, and R649W) at specific time points. The p110α/p85α-E601* variant was also tested but did not undergo changes in deuterium incorporation that met our threshold. **(D)** Schematic model depicting WT PI3Kα (p110α/p85α) and p85α variants in the presence of increasing levels of PDGFR pY. WT PI3Kα (p110α/p85α) SH2 domains are fully engaged by low levels of PDGFR pY, leading to breaking of inhibitory SH2 interfaces. The p85α variants (p110α/p85α-R590*, p110α/p85α-R649W) undergo similar or lesser changes as WT but only at the high concentrations of PDGFR pY.

## Discussion

Class IA PI3Ks are regulated through a complex network of inter and intra-protein interactions with its regulatory domains. Perturbations of the interaction between the p110 catalytic subunits and regulatory subunits through activating mutations is causative of multiple human diseases, including cancer (Fruman et al., 2017; Samuels et al., 2004), primary immunodeficiencies (Angulo et al., 2013; Dornan and Burke, 2018; Lucas et al., 2016), and developmental disorders (Goncalves and Cantley, 2018; Lindhurst et al., 2012; Rivière et al., 2012; Venot et al., 2018). Mutations mediate activation of PI3K activity through disruption of key inhibitory interfaces or through induction of allosteric conformational changes that mimic natural activation mechanisms of class IA PI3Ks (Burke et al., 2012; Dornan et al., 2017; Takeda et al., 2017). Class IA regulatory subunits can interact with all p110 catalytic subunits, so mutations in the genes that encode these subunits could be expected to perturb signalling through all p110 isotypes. Intriguingly, we have previously found that p110δ is specifically activated over p110α through an APDS2 splice site mutation (Δ434-475) in the iSH2 domain of *PIK3R1* (Dornan et al., 2017).

We were interested in further investigating the regulatory roles of disease linked mutations in the iSH2 and cSH2 domains of the p85α regulatory subunit encoded by *PIK3R1*, and any potential p110 isoform specific effects. The five class IA regulatory subunits all have a similar core domain architecture consisting of two N- and C-terminal SH2 domains connected through an anti-parallel coiled coil iSH2 domain. The iSH2 consists of three α-helices: α1, α2, and α3. Multiple truncations, insertions, deletions and point mutations have been discovered in the iSH2 domain in a variety of human diseases. These include a number of mutations/truncations found throughout N-terminus of the α1 helix as well as the C-terminal region of the iSH2 within the α2 and α3 helices (Cheung et al., 2011; Jaiswal et al., 2009; Jimenez et al., 1998; Urick et al., 2011). In addition, multiple mutations in the iSH2 domain have been discovered that are associated with the primary immunodeficiency APDS2 (Nunes-Santos et al., 2019), with the most common being a splice site mutation leading to deletion of exon 11 (residues 434-475), which is the N-terminal region of the α1 helix of the iSH2 (Deau et al., 2014; Lucas et al., 2014b). These previous mutations all lead to upregulated PI3K activity, however, mutations and truncations in both the iSH2 and cSH2 have been discovered that downregulate PI3K activity through decreased sensitivity to insulin signalling. These mutations cause SHORT syndrome (**S**hort stature, **h**yper-extensibility, hernia, **o**cular depression, **R**ieger anomaly, and **t**eething delay) (Chudasama et al., 2013; Dyment et al., 2013; Huang-Doran et al., 2016; Petrovski et al., 2016; Schroeder et al., 2014; Thauvin-Robinet et al., 2013), with the most common mutation disrupting the phosphotyrosine binding site in the cSH2 domain (R649W). We have characterised the molecular mechanism of activation of a C-terminal truncation of p85α (Q572*) and a point mutant in the cSH2 (R649W) that disrupts phosphorylated tyrosine binding leading to impaired insulin signalling. This has revealed novel isoform specific insight into the regulation of p110 catalytic activity by regulatory subunits, as well as how the p85α regulatory subunit mediates activation downstream of growth factor signalling.

To investigate the regulatory roles of the iSH2 domain, we characterised the p85α-Q572* oncogenic truncation, as well as truncations at the end of the α3 and α2 helices of the iSH2. The Q572* truncation in p85α led to activation of both the p110α and p110δ isoforms. This was especially striking for the p110α, where the Q572* mutant alone was ~40 fold more active than the wild type p110α/p85α complex in the basal state, and ~7 fold more active than this wild type upon stimulation by saturating concentrations of PDGFR pY. Conversely, p110δ in complex with the Q572* truncation showed a similar level of kinase activity to the wild type in the presence of saturating PDGFR pY. HDX-MS analysis on the p110α complex with Q572* revealed the molecular mechanism of activation, with this mutant leading to disruption of all inhibitory interfaces (ABD-kinase, C2-iSH2, nSH2-C2/helical/kinase). This is consistent with previous results showing that the combination of the activating point mutation in the C2 domain of p110α and the Q572* truncation does not lead to additive activation (Wu et al., 2009). However, the complex of Q572* with p110α does not represent a fully activated form, as kinase activity can still be stimulated by the presence of lipidated HRas similar to wild type p110α/p85α (Buckles et al., 2017; Siempelkamp et al., 2017).

The engineered and oncogenic C-terminal truncations (590* and 601* p85α, respectively) in complex with either p110α or p110δ led to greatly decreased activation downstream of a PDFGR derived bis-phosphorylated peptide. These experiments also revealed a partial role of the α3 helix of iSH2 in regulating lipid kinase activity for both p110α or p110δ, as this construct was able to be hyper-activated compared to the WT p85α, however, this required greatly increased levels of bis-phosphorylated PDGFR peptide to attain full activation. As both of these constructs do not contain the cSH2 domain, we hypothesized that perhaps these truncations were acting similar to previously described SHORT syndrome mutations. To test this, we generated p85α with a SHORT syndrome mutation (R649W) that disrupts the phosphopeptide binding site (Chudasama et al., 2013; Dyment et al., 2013; Huang-Doran et al., 2016; Thauvin-Robinet et al., 2013) in complex with both p110α and p110δ. We found that either loss of the cSH2 (601*) or mutation of the phosphopeptide binding site (R649W) in p85α led to similar decreased sensitivity to PDGFR pY, highlighting the importance of the cSH2 in proper PI3K signalling downstream of activated membrane receptors. The cSH2 has previously been shown to mediate high affinity binding of the p110/p85 complex to phosphorylated PDGFR (Felder et al., 1993; Panayotou et al., 1993; Songyang et al., 1993). To further verify this mechanism of activation we characterised the binding and activation by singly phosphorylated PDGFR phosphopeptides, and found that activation of wild type p110α/p85α by mono-phosphorylated PDGFR pY751 led to the same sensitivity to activation as p110α R649W-p85α with bis-phosphorylated PDGFR pY. To define the molecular basis for decreased sensitivity we carried out HDX-MS experiments with different doses of PDGFR pY for p110α bound to either p85α or R649W p85α. We found that for p110α/ p85α differences in HDX-MS were saturated at the lowest concentration (1 μM) of PDGFR pY, with no further changes at 20 μM. In the R649W complex we saw no changes in deuterium incorporation at 1 μM, with similar changes in exchange compared to the wild type at saturating levels (20 μM).

Mutations in *PIK3R1* are causative of multiple human diseases. This has led to great interest in defining the molecular mechanisms that mediate disease. Our study has shown that the Q572* oncogenic mutation of p85α leads to hyper-activation of p110α through disruption of all regulatory interfaces between p110 and p85. This hyper-activation is not observed for p110δ, revealing an intriguing isoform specific difference. Furthermore, our study reveals how SHORT mutations mediate downregulation of PI3K activity through greatly decreasing their sensitivity to bis-phosphorylated receptors and their adaptors. This study provides a framework for further study into the role of disease causing *PIK3R1* mutations in disease.

## STAR Methods

### Plasmid Generation

Plasmids harbouring WT p110α, p110δ, and p85α were previously described but in short, pFastBac 1 vector for expression through baculovirus expression vector protein expression systems in *Spodoptera frugiperda* (Sf9) cells contained the genes of interest (Dornan et al., 2017). The plasmids containing p110 isoforms also expressed N-terminal to the protein a 10X histidine tag, followed by a 2X Strep tag, followed by a Tobacco Etch Virus protease cleavage site. Single substitution mutations (R358A, R649W) and C-terminal truncations (Q572*, R590*, E601*) were generated using site-directed mutagenesis according to published commercial protocols (QuickChange Site-Directed Mutagenesis, Novagen). DNA oligonucleotides spanning the desired region and either containing the altered nucleotides (single substitutions) or lacking the truncated region were ordered (Sigma). PCR reactions were performed on the WT p85α and PCR purified (Q5 High-Fidelity 2X MasterMix, New England Bioscienes #M0492L; QiaQuick PCR Purification Kit, Qiagen #28104). Single colonies were grown overnight and purified using QIAprep Spin Miniprep Kit (Qiagen #27104). Plasmid identity was confirmed by sanger sequencing (Eurofins Genomics).

### Virus Generation and amplification

The pFastBac plasmids harbouring p110α, p110δ, p85α (WT and pathogenic variants) and HRas G12V were transformed into DH10MultiBac cells (MultiBac, Geneva Biotech) containing the MultiBac baculovirus viral genome (bacmid) and a helper plasmid expressing transposase to transpose the expression cassette harbouring the gene of interest into the baculovirus genome. Bacmids with successful incorporation of the expression cassette of pFastBac into the MultiBac viral genome were identified by blue-white screening and were purified from a single white colony using a standard isopropanol-ethanol extraction method. Briefly, colonies were grown overnight (~16 hours) in 3-5 mL 2xYT (BioBasic #SD7019). Cells were pelleted by centrifugation and the pellet was resuspended in 300 uL P1 Buffer (50 mM Tris-cl, pH 8.0, 10 mM EDTA, 100 ug/mL RNase A), chemically lysed by the addition of 300 uL Buffer P2 (1% sodium dodecyl sulfate (SDS) (W/V), 200 mM NaOH), and the lysis reaction was neutralized by addition of 400 uL Buffer N3 (3.0 M potassium acetate, pH 5.5). Following centrifugation at 21130 rcf and 4°C (Centrifuge 5424 R), the supernatant was separated and mixed with 800 uL isopropanol to precipitate the DNA out of solution. Further centrifugation at the same temperature and speed pelleted the Bacmid DNA, which was then washed with 500 uL 70% Ethanol three times. The Bacmid DNA pellet was then dried for 1 minute and re-suspened in 50 uL Buffer EB with light flicking (10 mM Tris-Cl, pH 8.5; All buffers from QIAprep Spin Miniprep Kit, Qiagen #27104).

Purified bacmid was then transfected into Sf9 cells. 2 mL of Sf9 cells between 0.3-0.5×10^6^ cells/mL were aliquoted into the wells of a 6-well plate and allowed to attach, creating a monolayer of cells at ~ 70-80% confluency. Transfection reactions were prepared by the addition of 2-10 ug of bacmid DNA to 100 uL 1xPBS and 12 uL polyethyleneimine (PEI) at 1 mg/mL (Polyethyleneimine “Max” MW 40.000, Polysciences #24765, USA) to 100 uL 1xPBS. The bacmid-PBS and the PEI-PBS solutions were mixed together, and the reaction occurred for 20-30 minutes before addition drop-by-drop to an Sf9 monolayer containing well. Transfections were allowed to proceed for 5-7 days before harvesting virus containing supernatant as a PI viral stock.

Viral stocks were amplified by adding P1 viral stock to suspension Sf9 cells between 1-2×10^6^ cells/mL at a 1/100 volume ratio. This amplification produces a P2 stage viral stock that can be used in final protein expression. The amplification proceeded for 4-5 days before harvesting, with cell shakings at 120 RPM in a 27°C shaker (New Brunswick). Harvesting of P2 viral stocks was carried out by centrifuging cell suspensions in 50 mL Falcon tubes at 2281 RCF (Beckman GS-15), collecting the supernatant in a fresh sterile tube, and adding 5-10% inactivated foetal bovine serum (FBS; VWR Canada #97068-085).

### Expression and Purification of all PI3K complexes

All PI3Kα and PI3Kδ complex variants were expressed and purified as previously described (Dornan et al., 2017). To express PI3K complexes, an optimised ratio of p110 subunit to p85α subunit (10 mL: 3 mL per 500 mL of insect cells) baculovirus was used to co-infect Sf9 cells between 1-2×10^6^ cells/mL. Co-infections were harvested between 42-72 hours by centrifugation at 1680 RCF (Eppendorf Centrifuge 5810 R) and pellets were washed with ice-cold 1xPBS before snap-freezing in liquid nitrogen. All PI3K variants were purified in an identical method by lysing cells and performing nickel affinity, streptavidin affinity, and size exclusion chromatography purification steps, with all steps carried out on ice or in a 4°C cold room.

Frozen Sf9 pellets were re-suspended in lysis buffer (20 mM Tris pH 8.0, 100 mM NaCl, 10 mM imidazole pH 8.0, 5% glycerol (v/v), 2 mM bME, protease inhibitor (Protease Inhibitor Cocktail Set III, Sigma)) and sonicated on ice for 1 minute 30 seconds (15s on, 15s off, level 4.0, Misonix sonicator 3000). Triton X-100 was added to the lysate at a concentration of 0.2% and centrifuged at 20,000 g for 45 minutes (Beckman Coulter Avanti J-25I, JA 25.50 rotor). The supernatant was then loaded onto a 5 mL HisTrap™ FF crude column (GE Healthcare #11000458) that had been equilibrated in NiNTA A buffer (20 mM Tris pH 8.0, 100 mM NaCl, 20 mM imidazole pH 8.0, 5% (v/v) glycerol, 2 mM bME). The column was washed with 20 mL of high salt NiNTA A buffer (20 mM Tris pH 8.0, 1 M NaCl, 20 mM imidazole pH 8.0, 5% (v/v) glycerol, 2 mM bME), 15 mL of NiNTA A buffer, 20 mL of 6% NiNTA B buffer (20 mM Tris pH 8.0, 100 mM NaCl, 500 mM imidazole pH 8.0, 5% (v/v) glycerol, 2 mM bME) before being eluted with 100% NiNTA B. The elution was loaded onto 2x tandem 1 mL StrepTrap™ HP columns (GE Healthcare #29048653) equilibrated in Hep A buffer (20 mM Tris pH 8.0, 100 mM NaCl, 5% (v/v) glycerol, 2 mM bME). The columns were washed with 2 mL of Hep A buffer before addition of a tobacco etch virus protease containing a stabilizing lipoyl domain. Tobacco etch virus cleavage proceeded overnight (~ 12–16 h) before elution of cleaved protein in 1–2 mL of Gel Filtration Buffer (20 mM HEPES pH 7.5, 150 mM NaCl, 0.5 mM tris(2-carboxyethyl) phosphine (TCEP)). Fractions were pooled and concentrated in a 50,000 MWCO Amicon concentrator (Millipore) to between 300-1000 uL. Concentrated protein was injected onto a Superdex™ 200 10/300 GL Increase size-exclusion column (GE Healthcare #28990944) equilibrated in Gel Filtration Buffer. Proteins were concentrated to a concentration between 0.5-4 mg/mL. Protein was aliquoted, snapfrozen in liquid nitrogen, and stored at -80 °C.

### Expression and Purification of Lipidated HRas G12V

Full-length HRas G12V was expressed and purified as previously described (Porfiri et al., 1995). The protein was expressed by infecting 500 mL of Sf9 cells with 5 mL of baculovirus. Cells were harvested after 55 hours of infection by centrifugation at 1680 RCF (Eppendorf Centrifuge 5810 R) following which cells were washed in ice-cold PBS and snap frozen in liquid nitrogen.

The frozen cell pellet was resuspended in lysis buffer (50 mM HEPES pH 7.5, 100 mM NaCl, 10 mM bME and protease inhibitor (Protease Inhibitor Cocktail Set III, Sigma)) and sonicated on ice for 1 minute 30 seconds (15s ON, 15s OFF, power level 4.0) on the Misonix sonicator 3000. Triton-X 114 was added to the lysate to a final concentration of 1%, mixed for 10 minutes at 4°C and centrifuged at 25,000 rpm for 45 minutes (Beckman Ti-45 rotor). The supernatant was warmed to 37°C for few minutes until it turned cloudy following which it was centrifuged at 11,000 rpm at room temperature for 10 minutes (Beckman JA-20 rotor) to separate the soluble and detergent-enriched phases. The soluble phase was removed, and Triton-X 114 was added to the detergent-enriched phase to a final concentration of 1%. This phase separation was performed for a total of 3 times. Imidazole pH 8.0 was added to the detergent phase to a final concentration of 15 mM and the mixture was incubated with Ni-NTA agarose beads (Qiagen) for 1 hour at 4°C. The beads were washed with 5 column volumes of Ras-NiNTA buffer A (20mM Tris pH 8.0, 100mM NaCl, 15mM imidazole pH 8.0, 10mM bME and 0.5% Sodium Cholate) and the protein was eluted with 2 column volumes of Ras-NiNTA buffer B (20mM Tris pH 8.0, 100mM NaCl, 250mM imidazole pH 8.0, 10mM bME and 0.5% Sodium Cholate). The protein was buffer exchanged to Ras-NiNTA buffer A using a 10,000 kDa MWCO Amicon concentrator, where protein was concentrated to ~1mL and topped up to 15 mL with Ras-NiNTA buffer A and this was repeated a total of 3 times. GTPgS was added in 2-fold molar excess relative to HRas along with 25 mM EDTA. After incubating for an hour at room temperature, the protein was buffer exchanged with phosphatase buffer (32 mM Tris pH 8.0, 200 mM Ammonium Sulphate, 0.1 mM ZnCl2, 10 mM bME and 0.5% Sodium Cholate). 1 unit of immobilized calf alkaline phosphatase (Sigma) was added per milligram of HRas along with 2-fold excess nucleotide and the mixture was incubated for 1 hour on ice. MgCl2 was added to a final concentration of 30 mM to lock the bound nucleotide. The immobilized phosphatase was removed using a 0.22-micron spin filter (EMD Millipore). The protein was concentrated to less than 1 mL and was injected onto a Superdex^TM^ 75 10/300 GL size exclusion column (GE Healthcare) equilibrated in gel filtration buffer (20 mM HEPES pH 7.7, 100 mM NaCl, 10 mM CHAPS, 1 mM MgCl2 and 2 mM TCEP). The protein was concentrated to 1 mg/mL using a 10,000 kDa MWCO Amicon concentrator, aliquoted, snap-frozen in liquid nitrogen and stored at -80°C.

### Lipid Vesicle Preparation

PI3K Lipid Kinase Vesicles - Lipid vesicles mimicking the composition of plasma membrane ((5% brain phosphatidylinositol 4,5-bisphosphate (PIP2), 30% brain phosphatidylserine (PS), 50% brain phosphatidylethanolamine (PE), 15% brain phosphatidylcholine (PC)) were prepared by combining lipid components dissolved in organic solvent, vigorously mixing, and evaporating the solvent under a stream of N2 gas while gently swirling the lipid mixture to ensure the production of an even lipid film layer. The lipid film was desiccated under vacuum for 30-60 minutes and resuspended at 1 mg/mL in lipid buffer (20 mM HEPES pH 7.5 (RT), 100 mM KCl, 0.5 mM EDTA). Vesicle solution was then sonicated for 10 minutes and subjected to three freeze-thaw cycles. Vesicles were then extruded 11 times through a 100-nm filter (T&T Scientific: TT-002-0010). Extruded vesicles were then snap-frozen in liquid nitrogen and stored at -80 °C.

Ras Activation Vesicles - Lipid vesicles containing 5% brain phosphatidylinositol 4,5-bisphosphate (PIP2), 20% brain phosphatidylserine (PS), 50% egg-yolk phosphatidylethanolamine (PE), 10% egg-yolk phosphatidylcholine (PC), 10% cholesterol and 5% egg-yolk sphingomyelin (SM) were prepared by mixing the lipids dissolved in organic solvent. The solvent was evaporated in a stream of nitrogen following which the lipid film was dessicated in a vacuum for 45 minutes. The lipids were resuspended in lipid buffer (20 mM HEPES pH 7.0, 100 mM NaCl and 10 % glycerol) and the solution was sonicated for 15 minutes. The vesicles were subjected to five freeze thaw cycles and extruded 11 times through a 100-nm filter (T&T Scientfic: TT-002-0010). The extruded vesicles were sonicated again for 5 minutes, aliquoted and stored at -80°C.

### PDGFR phosphopeptide (pY)

All peptides were custom ordered from New England Peptide (NEP). Three peptides in total were ordered. The initial peptide spans residues 735-767 of the platelet derived growth factor receptor (PDGFR) and is phosphorylated at residues 740 (pY740) and 751 (pY751). The other two peptides were prepared with the same residues, differing in their phosphorylation. Each single peptide is denoted by its phosphorylated residue (pY740 or pY751).

### Lipid Kinase Assays

Lipid kinase activity was determined using the TRANSCREENER^®^ ADP2 Fluorescence Intensity (FI) Assay (Bellbrook Labs: 3013-1 K), which monitors the hydrolysis of ATP. For PI3K and PDGFR pY activation of PI3K assays, vesicles were used at a final concentration of 0.45 mg/mL, and ATP was used at a final concentration of 100 μM. Protein (4X final concentration) was incubated with PDGFR pY (4X final concentration) or a blank solution for 6-9 minutes before starting the reaction by the addition of 2 μL of 2X start mixture (ATP, lipid vesicles) to 2 μL of 2X protein or protein/PDGFR pY mixture, and allowed to proceed for 60 min at 27 °C. Reactions were stopped by the addition of 4 μL of 2 X stop and detect mixture (1 × Stop and Detect Buffer, 8 nM ADP Alexa Fluor 594 Tracer, 93.7 μg/mL ADP2 Antibody-IRDye QC-1) and allowed to incubate for a minimum of 45 min. The fluorescence intensity was determined using a SpectraMax M5 plate reader with an excitation at 590 nm and emission at 620 nm. Relative fluorescence data obtained was normalised to the assay window (100% ADP – 0% ADP), and the %ATP turnover was calculated based on an ATP standard curve prepared for the concentration of ATP used in the assay. The %ATP turnover was then used to calculate the specific activity (nmol ADP/mg of enzyme*min). Comparisons between conditions were tested using the standard students t-test and p values <0.01 were considered significant.

For Ras Activation Assays, 2X start mixture has 3 μM HRas G12V GTPgS incubated with vesicles and ATP for 5 minutes at room temperature. Lipid vesicles were used at a final concentration of 0.5 mg/mL and ATP was used at a final concentration of 100 μM. The reaction was started by adding 2 μL of 2X start mixture to 2 μL of 2X protein and allowed to proceed for 60 min at 20 °C. Following this, the reaction was stopped, the intensity was measured, and specific activity calculated as described above.

### Hydrogen Deuterium eXchange

#### Sample Preparation

All HDX experiments were conducted in 20 μL reactions with a final concentration of 500 nM (p85α-Q572* experiment; Fig. 2) or 660 nM (PDGFR pY p85α mutation/truncations experiment; Fig. 4) for WT PI3Kα and PI3Kα deletions. In the p85α-Q572* experiment, WT PI3Kα and p85α-Q572* truncation PI3Kα were only measured in the basal condition. For the PDGFR pY experiment, three conditions were tested: PI3Kα alone, PI3Kα in the presence of low concentration PDGFR phosphopeptide (1 μM pY), and PI3K in the presence of high concentration PDGFR phosphopeptide (20 μM pY). Deuterium exchange was initiated by adding 16 μL of deuterated buffer (10 mM HEPES pH 7.5, 100 mM NaCl, 96% (v/v) D2O) to 4 uL of protein or protein/PDGFR pY mixture. Exchange was carried out for three time points (3s, 30s, and 300s at 23°C) and terminated by the addition of 50 μL ice-cold quench buffer (0.8 M guanidine-HCl, 1.2% formic acid). After quenching, samples were immediately frozen in liquid nitrogen and stored at -80°C. All experiments were carried out in triplicate.

#### Protein Digestion and MS/MS Data Collection

Protein samples were rapidly thawed and injected onto an integrated fluidics system containing a HDx-3 PAL liquid handling robot and climate-controlled chromatography system (LEAP Technologies), a Dionex Ultimate 3000 UHPLC system, as well as an Impact HD QTOF Mass spectrometer (Bruker). The protein was run over two immobilized pepsin columns (Applied Biosystems; Poroszyme™ Immobilized Pepsin Cartridge, 2.1 mm x 30 mm; Thermo-Fisher 2-3131-00; at 10°C and 2°C respectively) at 200 μL/min for 3 minutes. The resulting peptides were collected and desalted on a C18 trap column (Acquity UPLC^®^ BEH^TM^ C18 1.7μm column (2.1 x 5 mm); Waters 186002350). The trap was subsequently eluted in line with a C18 reverse-phase separation column (Acquity 1.7 μm particle, 100 × 1 mm^2^ C18 UPLC column, Waters 186002352), using a gradient of 5-36% B (Buffer A 0.1% formic acid; Buffer B 100% acetonitrile) over 16 minutes. Mass spectrometry experiments were performed on an Impact II QTOF (Bruker) acquiring over a mass range from 150 to 2200 *m/z* using an electrospray ionization source operated at a temperature of 200 °C and a spray voltage of 4.5 kV.

#### Peptide Identification

Peptides were identified using data-dependent acquisition following tandem MS/MS experiments (0.5 s precursor scan from 150-2000 m/z; twelve 0.25 s fragment scans from 150-2000 m/z). MS/MS datasets were analyzed using PEAKS7 (PEAKS), and a false discovery rate was set at 1% using a database of purified proteins and known contaminants.

#### Mass Analysis of Peptide Centroids and Measurement of Deuterium Incorporation

HD-Examiner Software (Sierra Analytics) was used to automatically calculate the level of deuterium incorporation into each peptide. All peptides were manually inspected for correct charge state, correct retention time, appropriate selection of isotopic distribution, etc. Deuteration levels were calculated using the centroid of the experimental isotope clusters. Results are presented as relative levels of deuterium incorporation and the only control for back exchange was the level of deuterium present in the buffer (76.8% for all experiments). Changes in any peptide at any time point greater than specified cut-offs (both 7% and 0.4 Da between conditions for the Q572* mutant versus wild-type and both 7% and 0.4 Da between conditions for the PDGFR pY dose experiment) and with an unpaired t-test value of p<0.05 was considered significant. The full set of HDX-MS data is presented in Source Data 1, with all HDX-MS analysis statistics shown in Table S1 as suggested by the guidelines set out in (Masson et al., 2019).

#### Isothermal Titration Calorimetry

WT PI3Kα and R649W PI3Kα were purified with a final buffer of 20 mM Hepes pH 7.5, 150 mM NaCl and 1 mM TCEP. Peptide stocks were diluted in the same ITC buffer as above. All ITC experiments were carried out at 20 °C on a MicroCal iTC200 instrument (GE Healthcare). The sample cell contained WT p110α/p85α (7.6 μM) or p110α/ R649W p85α (11 μM), and peptide (46-132 μM) was added in 19 injections of 2 μL each. Data was processed using Origin software (MicroCal) and the dissociation constants (Kd) were determined using a one-site model. Figures are of a single experiment, and are representative of at least two independent experiments.

## Acknowledgements

This work was supported by a Cancer Research Society Operating Grant CRS-22641, a Canadian Institutes of Health Research New Investigator award, and a Michael Smith Foundation for Health Research Scholar Award 17686.

## Author Contributions

All biochemical experiments, protein purification, and molecular biology was carried out by GLD. HRas activation assays were carried out by MKR. HDX-MS experiments were carried out by GLD and JTBS. ITC experiments were carried out and analyzed by CJP and MJB. Data was analysed by GLD, JTBS, and JEB. Experiments were conceived and designed by JEB and GLD. The manuscript was written by JEB and GLD with input from all authors.

