## Supplemental data for "Defining how oncogenic and developmental mutations of *PIK3R1* alter the regulation of class IA phosphoinositide 3-kinases"

### Supplemental Figures and Figure Legends

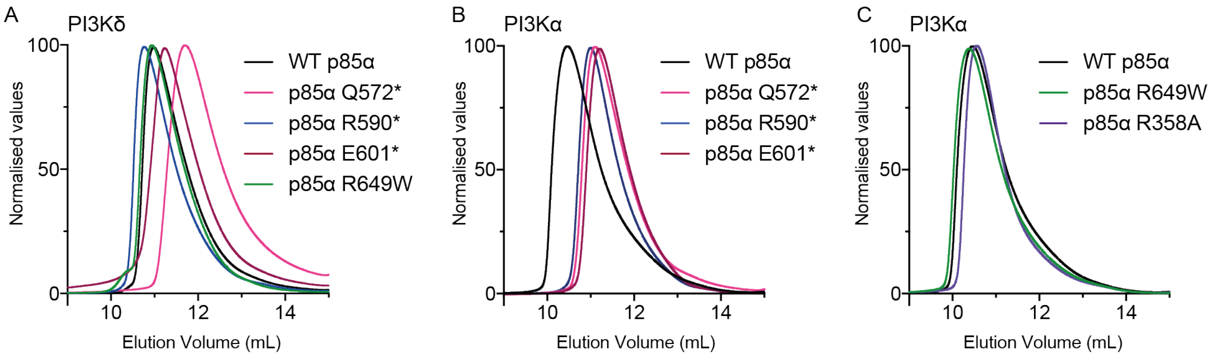


**Figure S1. Size exclusion chromatography (SEC) traces of WT and mutant proteins.** All SEC was carried out using an S200 10/300 GL SEC column (GE Healthcare) under the same buffer conditions as described in the methods. The traces for **(A)** PI3Kδ complexes composed of p110δ and deletion or point mutation variants of p85α. (**B**) PI3Kα complexes composed of p110α and deletion variants of p85α. (**C**) PI3Kα complexes composed of p110α and p85α mutants of the nSH2 or cSH2 FLVR motifs.

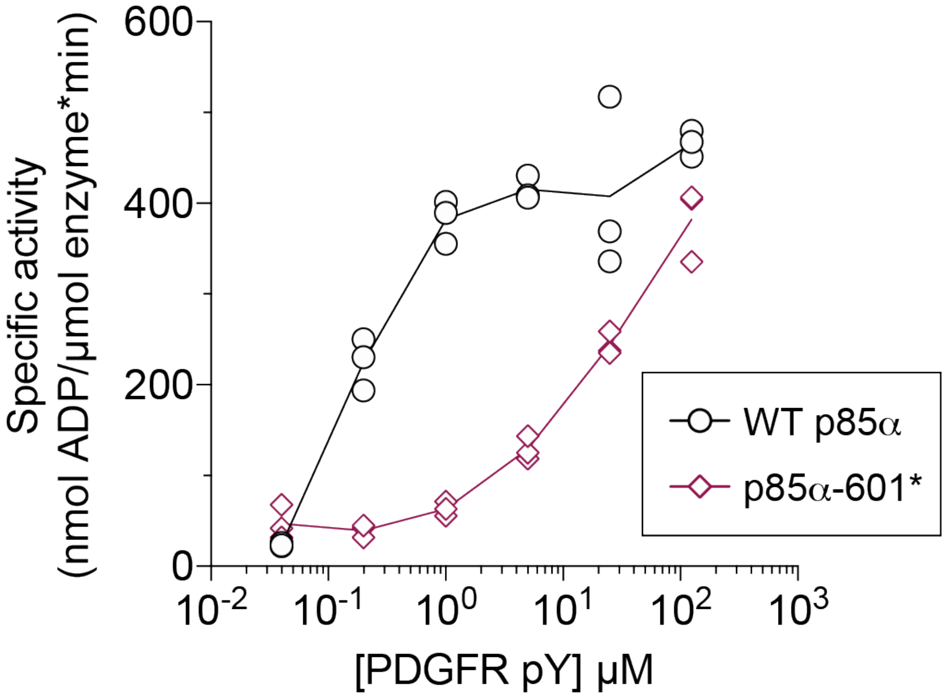


**Figure S2. C-terminal truncation of p85α leads to decreased sensitivity to PDGFR stimulation in p110δ.** Dose response of bis-phosphorylated PDGFR phosphopeptide concentration of the wild-type and the C-terminal truncation E601*. Assays measured the production of ADP in the presence of 50–1100 nM of enzyme, 100 μM ATP, and PM mimic vesicles containing 5% PIP2. Kinase assays were performed in triplicate (error shown as SD; n = 3-6).

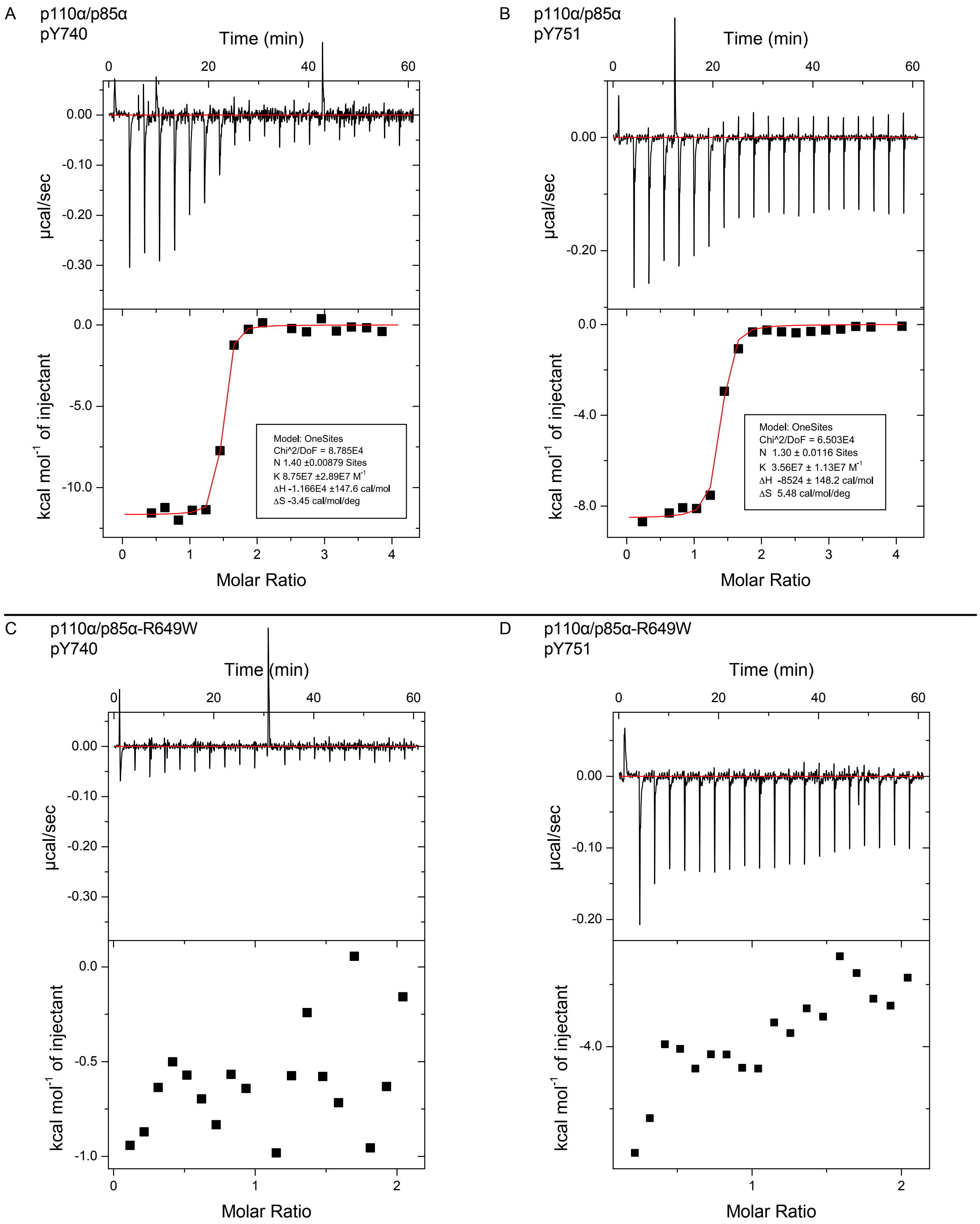


**Figure S3.** PI3K binds two phosphopeptide motifs simultaneously, requires functional SH2 domain. Top) Representative ITC binding isotherms of p110α/p85α following titration of single PDGFR phosphopeptide with either A) the pY740 site or B) the pY751 site. Bottom) Representative ITC binding isotherms of the SHORT mutant p110α/p85α-R649W following titration of single PDGFR phosphopeptide with either C) the pY740 site or D) the pY751 site.

| Data Set (Fig 2) | P110a/p85a | P110a/ Q572*p85a |
|---|---|---|
| HDX reaction details | %$D_2O$=76.8%<br>$pH_{(read)}$= 7.5<br>Temp= 20°C | %$D_2O$=76.8%<br>$pH_{(read)}$= 7.5<br>Temp= 20°C |
| HDX time course | 3s, 30s, 300s | 3s, 30s, 300s |
| HDX controls | N/A | N/A |
| Back-exchange | Corrected based on %$D_2O$ | Corrected based on %$D_2O$ |
| Number of peptides | P110a= 169<br>P85a=112 | P110a= 169<br>P85a= 91 |
| Sequence coverage | P110a= 92.9%<br>P85a= 91.6% | P110a= 92.9%<br>P85a= 91.6% |
| Average peptide length/ redundancy | P110a= 12.6 P85a= 14.4<br>Redundancy P110a= 2.0<br>Redundancy P85= 1.8 | P110a= 12.6 P85a= 13.9<br>Redundancy P110a= 2.0<br>Redundancy P85a= 1.8 |
| Replicates | 3 | 3 (only 2 for 30s time point) |
| Repeatability | Average StDev = 0.6% | Average StDev = 0.8% |
| Significant differences in HDX | >7% and >0.4 Da and unpaired t-test <0.05 | >7% and >0.4 Da and unpaired t-test <0.05 |

| Data Set (Fig 4) | P110a/p85a | P110a/p85a +1 uM pY | P110a/p85a+20 uM pY | P110a/p85a-R590* | P110a/p85a-R590* +1 uM pY | P110a/p85a-R590*+20 uM pY |
|---|---|---|---|---|---|---|
| HDX reaction details | %$D_2O$=76.8%<br>$pH_{(read)}$= 7.5<br>Temp= 20°C | %$D_2O$=76.8%<br>$pH_{(read)}$= 7.5<br>Temp= 20°C | %$D_2O$=76.8%<br>$pH_{(read)}$= 7.5<br>Temp= 20°C | %$D_2O$=76.8%<br>$pH_{(read)}$= 7.5<br>Temp= 20°C | %$D_2O$=76.8%<br>$pH_{(read)}$= 7.5<br>Temp= 20°C | %$D_2O$=76.8%<br>$pH_{(read)}$= 7.5<br>Temp= 20°C |
| HDX time course | 3s, 30s, 300s | 3s, 30s, 300s | 3s, 30s, 300s | 3s, 30s, 300s | 3s, 30s, 300s | 3s, 30s, 300s |
| HDX controls | N/A | N/A | N/A | N/A | N/A | N/A |
| Back-exchange | Corrected based on %$D_2O$ | Corrected based on %$D_2O$ | Corrected based on %$D_2O$ | Corrected based on %$D_2O$ | Corrected based on %$D_2O$ | Corrected based on %$D_2O$ |
| Number of peptides | P110a= 143<br>P85a=112 | P110a= 143<br>P85a=112 | P110a= 138<br>P85a=112 | P110a= 143<br>P85a=94 | P110a= 143<br>P85a=94 | P110a= 138<br>P85a=94 |
| Sequence coverage | P110a= 85.2%<br>P85a= 93.0% | P110a= 85.2%<br>P85a= 93.0% | P110a= 84.4%<br>P85a= 93.0% | P110a= 85.2%<br>P85a= 93.0% | P110a= 85.2%<br>P85a= 93.0% | P110a= 84.4%<br>P85a= 93.0% |
| Average peptide length/ redundancy | P110a= 12.3<br>P85a= 13.7<br>Redundancy P110a= 1.7<br>Redundancy P85= 2.1 | P110a= 12.3<br>P85a= 13.7<br>Redundancy P110a= 1.7<br>Redundancy P85= 2.1 | P110a= 12.4<br>P85a= 13.7<br>Redundancy P110a= 1.6<br>Redundancy P85= 2.1 | P110a= 12.3<br>P85a= 13.5<br>Redundancy P110a= 1.7<br>Redundancy P85= 2.1 | P110a= 12.3<br>P85a= 13.5<br>Redundancy P110a= 1.7<br>Redundancy P85= 2.1 | P110a= 12.4<br>P85a= 13.5<br>Redundancy P110a= 1.6<br>Redundancy P85= 2.1 |
| Replicates | 3 | 3 | 3 (only 2 for 3s and 300s time points) | 3 | 3 | 3 |
| Repeatability | Average StDev = 0.6% | Average StDev = 0.6% | Average StDev = 0.6% | Average StDev = 0.6% | Average StDev = 0.6% | Average StDev = 0.6% |
| Significant differences in HDX | >7% and >0.4 Da and unpaired t-test <0.05 | >7% and >0.4 Da and unpaired t-test <0.05 | >7% and >0.4 Da and unpaired t-test <0.05 | >7% and >0.4 Da and unpaired t-test <0.05 | >7% and >0.4 Da and unpaired t-test <0.05 | >7% and >0.4 Da and unpaired t-test <0.05 |

| Data Set (Fig 4) | P110a/p85a-R649W | P110a/p85a-R649W+1 uM pY | P110a/p85a-R649W+20 uM pY |
|---|---|---|---|
| HDX reaction details | %$D_2O$=76.8%<br>$pH_{(read)}$= 7.5<br>Temp= 20°C | %$D_2O$=76.8%<br>$pH_{(read)}$= 7.5<br>Temp= 20°C | %$D_2O$=76.8%<br>$pH_{(read)}$= 7.5<br>Temp= 20°C |
| HDX time course | 3s, 30s, 300s | 3s, 30s, 300s | 3s, 30s, 300s |
| HDX controls | N/A | N/A | N/A |
| Back-exchange | Corrected based on %$D_2O$ | Corrected based on %$D_2O$ | Corrected based on %$D_2O$ |
| Number of peptides | P110a= 143<br>P85a=112 | P110a= 143<br>P85a=112 | P110a= 138<br>P85a=112 |
| Sequence coverage | P110a= 85.2%<br>P85a= 93.0% | P110a= 85.2%<br>P85a= 93.0% | P110a= 84.4%<br>P85a= 93.0% |
| Average peptide length/ redundancy | P110a= 12.3<br>P85a= 13.7<br>Redundancy P110a= 1.7<br>Redundancy P85= 2.1 | P110a= 12.3<br>P85a= 13.7<br>Redundancy P110a= 1.7<br>Redundancy P85= 2.1 | P110a= 12.4<br>P85a= 13.7<br>Redundancy P110a= 1.6<br>Redundancy P85= 2.1 |
| Replicates | 3 | 3 | 3 |
| Repeatability | Average StDev = 0.6% | Average StDev = 0.6% | Average StDev = 0.6% |
| Significant differences in HDX | >7% and >0.4 Da and unpaired t-test <0.05 | >7% and >0.4 Da and unpaired t-test <0.05 | >7% and >0.4 Da and unpaired t-test <0.05 |

**Table S1**. Summary of HDX-MS parameters

| **Phosphopeptide** | **Molar Ratio** | $K_d$ (nM) | **ΔH** (kcal/mol) | **-TΔS** (kcal/mol) |
|---|---|---|---|---|
| Peptide 1 (pY740) | 1.49 ± 0.12 | 15.7 ± 6.1 | -11.7 ± 0.1 | 1.2 ± 0.3 |
| Peptide 2 (pY751) | 1.30 ± 0.01 | 28.0 ± 8.9 | -8.5 ± 0.1 | 1.6 |

**Table S2**. Summary of ITC thermodynamic parameters. All experiments performed in duplicate. Thermodynamic parameters fitted using Origin software (Microcal).
